# Porcine and human aortic valve endothelial and interstitial cell isolation and characterization

**DOI:** 10.1101/2022.12.01.518669

**Authors:** D. Nehl, PR. Goody, K. Maus, A. Pfeifer, E. Aikawa, F. Bakthiary, S. Zimmer, G. Nickenig, F. Jansen, MR. Hosen

## Abstract

**Background:** Calcific aortic valve stenosis is defined by pathological changes in the aortic valve and their predominant cell types: valvular interstitial (VICs) and endothelial cells (VECs). Understanding the cellular and molecular mechanisms of this disease is a prerequisite to identify potential pharmacological treatment strategies. In this study, we present a unique aortic valve cell isolation technique to acquire specific human and porcine cell populations and compared VICs and VECs of these species with each other for the first time.

**Methods and Results:** Aortic valve cells were isolated from human explants from patients undergoing surgical aortic valve replacement or porcine valvular tissue. Pure VEC and VIC populations could be verified by gene expression analysis and immunofluorescence staining showing a highly significant upregulation of endothelial markers in VECs and mesenchymal markers in VICs, respectively. Further analysis and comparison of cells in *in vitro* experiments revealed that endothelial-to-mesenchymal transition could be induced in hVECs, leading to significant increase of mesenchymal markers. *In vitro* calcification experiments of VICs induced by osteogenic medium or pro-calcifying medium demonstrated a pronounced calcification marker expression and visible calcific deposition in Alizarin red staining in both species.

**Conclusion:** This study aims to initiate a first step towards standardization of a reproducible isolation technique for pure human and porcine VEC and VIC populations. Comparison of human and porcine aortic valve cells demonstrated that porcine cells might serve as an alternative cellular model system, in settings, where human tissues are difficult to obtain.

**Statements and Declarations:** The authors declare no relevant financial or non-financial interests to disclose.

## Introduction

Aortic valve stenosis (AVS) due to calcific aortic valve disease (CAVD) is one of the most common heart valve diseases worldwide, with a 2-year mortality greater than 50% when symptomatic [1–3]. CAVD is characterized by fibrous thickening and calcification of the aortic valve cusps, leading to narrowing of the valve orifice. Originally, this disease was described as passive and degenerative but it is now well established that CAVD is a cell-mediated, actively regulated inflammatory disease [4–7]. Currently, there are no approved medical therapies to prevent or reverse calcification of the aortic valve. The only treatment options are open heart surgery with aortic valve replacement (SAVR) or transcatheter aortic valve implantation (TAVR) [8]. It is of upmost importance to better understand the mechanisms underlying this disease in order to identify feasible pharmaceutical and non-surgical/interventional therapies.

In healthy humans, the tricuspid aortic valve (AV) consists of three cusps, which are covered by valvular endothelial cells (VECs) on the aortic and ventricular sides. The interstitium of the valve is composed of three main layers with ubiquitously distributed valvular interstitial cells (VICs). During the pathogenesis of AVS, different stimuli, including shear stress and mechanical strain, trigger VEC dysfunction, resulting in lipoprotein/phospholipid infiltration and oxidation [9–11]. VECs lose their protective and regulative properties and can transdifferentiate into myofibroblast-like VICs, a process referred to as endothelial to mesenchymal transition (EndMT) [12–14]. Disruption of the endothelial layer also enables infiltration of immune cells into the AV, promoting the release of pro-inflammatory stimuli, and driving osteogenic differentiation of VICs [9, 15]. This osteogenic differentiation of VICs, from a fibroblastic to an osteogenic phenotype, induces the formation of calcium nodules, thus leading to AV calcification [9, 16]. It is pivotal to further investigate these molecular and cellular mechanisms of AV cell populations involved in CAVD. However, isolation and cultivation of VECs and VICs is technically and methodologically highly demanding. Here, we offer a novel AV cell isolation method that can be applied in all cell culture laboratories.

A variety of organisms can be used for such cell culture experiments, including human and pigs [17–20]. In general, human material is advantageous over animal material when studying potential therapeutic targets involved in disease pathogenesis in humans. While tissue from patients with CAVD can be used for genetic screening, cells from non-stenotic and healthy patients are the most useful and authentic platform to study disease progressions (e.g., induction of osteoblastic differentiation and EndMT). Calcified cusps from diseased human AVs can be obtained from patients undergoing AV replacement surgery due to severe AVS. Control tissues are more difficult to obtain, since non-calcified aortic valves are less often explanted (e.g. in the setting of aortic root or ascending aorta dilatation/aneurysm, when aortic regurgitation occurs as a consequence). It is therefore of importance to investigate whether other organisms, which are genetically comparable to humans, can be used as tissue donors for cell isolation. Undoubtedly, acquiring high-quality control samples from healthy or non-stenotic AVs is challenging. In addition to the most obvious problems, individual patients naturally display a high variability in age, sex, congenital, pre-existing, and concomitant diseases [21]. Thus, the degrees of calcification patterns differ immensely between patients and with respect to sex. Due to these difficulties, the most frequently used AVs tissues for cell isolation are obtained from large animals which resemble humans genetically, such as pigs [22–24]. An advantage of porcine tissue over human material is that healthy tissue can be obtained more easily from abattoirs or animal research facilities. The hearts and AVs are healthy without being affected by other diseases, and the cells can therefore be isolated and expanded more efficiently *in vitro*. The size and the substantial amount of tissue in porcine AVs makes it easier to study the cusps’ mechanical properties or to isolate cells. Moreover, the whole heart can be used for research, whereas in humans usually only one valve is explanted [25].

Although porcine valves appear to be anatomically and genetically similar to humans, there could be differences in specific molecular pathways. To date, no direct comparison of human and porcine VECs and VICs has been undertaken [26]. In this study, we undertook a major comparison of pig and human AV cells in order to identify a more easily available alternative to human material for cell culture experiments. Cells were isolated in a cell population-specific manner and characterized further. Cell culture experiments, such as *in vitro* EndMT and calcification induction, were performed in human and porcine cells to analyze and compare their response to these external stimuli.

## Methods

### Human and porcine aortic valve cell isolation

Human AV samples were obtained from patients undergoing AV replacement due to symptomatic AVS, or aortic regurgitation due to aortic root or ascending aorta dilatation, at Heart Center Bonn, University Hospital Bonn, Germany. The study protocol was approved by the ethics committee of the University Hospital Bonn (approval number AZ 078/17). All patients signed informed consent and the study was performed in concordance with the Declaration of Helsinki and International Conference on Harmonization of Good Clinical Practice [27]. After explantation, AV cusps were placed in 1x PBS on ice and transported to the laboratory immediately for cell isolation. Porcine AVs were gathered from an abattoir in Euskirchen, Germany. Freshly slaughtered pigs served as donors and, hearts were directly rinsed with NaCl and placed in 500 mL cold 1 x PBS on ice and supplemented with 2500 U heparin-sodium (B. Brown). Three AV cusps were excised from the aortic root. Directly after explantation, human aortic valve endothelial cells (hVEC), human aortic valve interstitial cells (hVICs), porcine aortic valve endothelial cells (pVECs) and porcine aortic valve interstitial cells (pVICs) were isolated from the three cusps starting with an incubation in 600 U/mL collagenase II in VIC primary medium (DMEM (Thermo Fisher Scientific, 21885108) supplemented with 10% FBS (Gibco, #A3160802), 1% penicillin/ streptomycin (pen/ strep) (Merck, #516106), and 44 mM NaHCO_3_ (AppliChem, #A1353)) for 10 minutes at 37 °C with gentle agitation. Cusps were placed in a petri dish filled with VEC medium (EGM™-2 MV Microvascular Endothelial Cell Growth Medium-2 BulletKit™ (Lonza, #C-3202)) and VECs were scraped into the medium using a sterile scalpel. Cells were washed twice with VEC medium by centrifugation for 15 minutes at 1000 rpm and filtered through a 100 μm cell strainer. To ensure a pure VEC suspension, cells were magnetically labeled with CD105 Magnetic Beats (Miltenyi Biotech, #130-051-201) and a MACS separation was performed according to the manufacturer’s protocol. VECs were seeded on fibronectin-coated (Sigma-Aldrich, #F0895) (5 μg/cm^2^) T25 flasks. To isolate VICs, the remaining valve tissue was minced and further digested using 600 U/mL collagenase II in VIC primary medium for 20h at 37 °C with gentle agitation. The cell suspension was then filtered (100 μm) and centrifuged twice for 15 minutes at 1000 rpm. VICs were seeded on collagen-coated (5 μg/cm^2^) T75 flasks.

### Cell culture

All cells were split with a detach kit from Lonza (Lonza, #CC-5034) and cultured in sterile incubators at 37°C, 5% (v/v) CO_2_ and 95% humidity. hVECs and pVECs were cultured in EGM™-2 MV Microvascular Endothelial Cell Growth Medium-2 BulletKit™ (Lonza, #CC-3202). hVICs and pVICs were cultured in VIC medium, consisting of DMEM (Thermo Fisher Scientific, 21885108), 10% FBS (Gibco, #A3160802) and 1% penicillin/streptomycin (pen/strep) (Merck, #516106). All experiments were performed in the fifth or sixth passage at 70% confluency on 12-or 24-well plates.

### Migration assay

Cell migration was analyzed by scratch wound assay [28] on a 24-well plate. hVECs and pVECs were incubated with starving medium, VEC medium without FBS, for 18 hours and subsequently, a scratch was performed using a sterile 200 μl pipet tip. The medium was changed to normal VEC medium, and the scratch wound closure documented every 2 hours by taking pictures with a Zeiss Axiovert 200M Microscope. The remaining cell-free area under the scratch was analyzed by Zen Lite software (Zeiss).

### *In vitro* EndMT induction

*In vitro* endothelial-to-mesenchymal transition (EndMT) was induced in hVECs and pVECs on a 12-well plate using EGM™-2 MV Microvascular Endothelial Cell Growth Medium-2 BulletKit™ (Lonza, #CC-3202), without VEGF, GA-1000 and hydrocortisone, but supplemented with 30ng/μl tumor necrosis factor alpha (TNFα; R&D systems, #210-TA-020) for 7d. EndMT induction was verified by gene expression analysis by RT-qPCR.

### *In vitro* calcification model

For *in vitro* calcification, hVICs and pVICs were seeded onto 12-well plates and cultured with DMEM (Thermo Fisher Scientific, #21885108), 5% FBS (Gibco, #A3160802) and 1% penicillin/ streptomycin (pen/ strep) (Merck, #516106), osteogenic medium (OM) or pro-calcifying medium (PCM). OM consisted of DMEM with 5% FBS, 1% pen/strep, 10 nmol/L β-glycerophosphate (Sigma-Aldrich, #G9422), 10 mmol/L dexamethasone (Sigma-Aldrich, #D4902) and 50 μg/mL L-ascorbic acid (Carl Roth, #3525.2). PCM is composed of DMEM with 5% FBS, 1% pen/ strep, 2 mmol/L sodium dihydrogen phosphate (NaH_2_PO_4_; Merck, #71507) and 50 μg/mL L-ascorbic acid (Carl Roth, #3525.2). Cells were incubated for 7 days to analyze gene expression by RT-qPCR and 21 days for calcium deposition staining with 2% alizarin red (Sigma-Aldrich, #A5533). For this purpose, cells were fixed with 4% formaldehyde for 15 minutes and washed twice with distilled water. Calcium nodules were stained with Alizarin red for 15 minutes at room temperature, followed by two further washing steps.

### RNA isolation and real time PCR-based gene expression profiling

RNA of cultured VICs and VECs was extracted using TRIzol reagent (Thermo Fisher Scientific, #15596018) and the following cDNA synthesis was performed with High-Capacity cDNA Reverse Transcription Kit (Thermo Fisher Scientific, # 43-688-13), both according to the manufacturer’s instructions. RT PCR-based gene expression profiling was conducted with TaqMan™ Gene expression master mix (Thermo Fisher Scientific, #4370074) and the following human TaqMan™ primers (Thermo Fisher Scientific): *GAPDH* (Hs02758991_g1), *α-SMA* (Hs00426835_g1), *vWF* (Hs01109446_m1), *PECAM-1* (Hs01065279_m1), *VIM* (Hs00418522_m1). The following porcine TaqMan™ Primers were used: *GAPDH* (Ss03375629_u1), *α-SMA* (Ss04245588_m1), *vWF* (Ss04322692_m1), *PECAM-1* (Ss04322692_m1), *VIM* (Ss04330801_gH), *ALPL* (Ss06879568_m1), *BMP2* (Ss03373798_g1), *BGLAP* (Ss03373655_s1), *SPP1* (Ss03391321_m1). ΔCT values were calculated with Microsoft Excel (Microsoft Corp., Redmond, WA, USA) normalizing to *GAPDH*, and significance tested and visualized with GraphPad Prism (GraphPad Software, Inc., San Diego, CA, USA).

### Immunofluorescence staining

Cells were fixed in 24-well plates with glass coverslips for 30 minutes with 4% formaldehyde. After three further washing steps with 1x PBS, cells were permeabilized using 0.25% Triton-X in PBS for 10 minutes and then washed again three times with 1x PBS. Cells were blocked with 1% BSA-glycine in PBS with 0.1% Tween for 30 minutes and then incubated for 2 hours with the primary antibodies on a shaker. For hVECs and hVICs alpha smooth muscle actin (α-SMA) (Abcam, #ab7817), vimentin (VIM) (Abcam, #8978), von Willebrand Factor (vWF) (Abcam, #ab6994) and platelet adhesion molecule-1 (PECAM-1) (Abcam, #28364) were used. For pVECs and pVICs, the vWF antibody was replaced by vWF (Bio-Rad, #AHP062T). Cells were washed three times with 1x PBS followed by incubation with the secondary antibody on the shaker for an hour: goat IgG anti-mouse IgG Cy3 (Dianova, #115-165-146), goat IgG anti-rabbit IgG Cy2 (Dianova, #111-225-144) or donkey IgG anti-sheep IgG Cy2 (Dianova, #712-225-147). Again, cells were washed three times with 1x PBS and embedded with Vectashield® Antifade Mounting Medium with DAPI (Vectashield, #H-1200-10). Fluorescence microscopy was performed with an Axio Observer Inverted Microscope (Carl Zeiss, Jena, Germany).

## Results

### Cell isolation leads to pure hVEC and hVIC populations

MACS-sorted hVEC and hVIC were characterized by RT-qPCR to quantify cell specific markers. Our quantification revealed a significant upregulation in endothelial marker expression in hVECs, such as von Willebrand factor (*vWF*) and platelet endothelial adhesion molecule-1 (*PECAM-1*) compared to hVICs. Myofibroblastic markers, alpha smooth muscle actin (*α-SMA*) and vimentin (*VIM*), were downregulated (Figure 1b). Immunofluorescence staining demonstrated an increase of intracellularly localized vWF and an expression of the endothelial marker, PECAM-1, whereas no expression of α-SMA and VIM was observed, further providing evidence that isolated cells were hVEC (Figure 1c). Analysis of hVEC migration capacity via a scratch wound closure revealed only 15% scratch width after 8 hours (Figure 1d), suggesting these cells maintain endothelial migration capacities, a further confirmation of pure hVEC population. To further assess if hVECs were able to undergo EndMT *in vitro*, cells were incubated with 30 ng/μl TNFα for 7 days. hVECs displayed increased expression of mesenchymal markers, such as *α-SMA* and *VIM*, and decreased expression of endothelial markers, namely *vWF* and *PECAM-1* (Figure 1e), demonstrating hVEC’s mesenchymal transdifferentiation capacity. These data suggest that elevated mesenchymal marker expression represents an effective EndMT induction.

**Figure 1:**
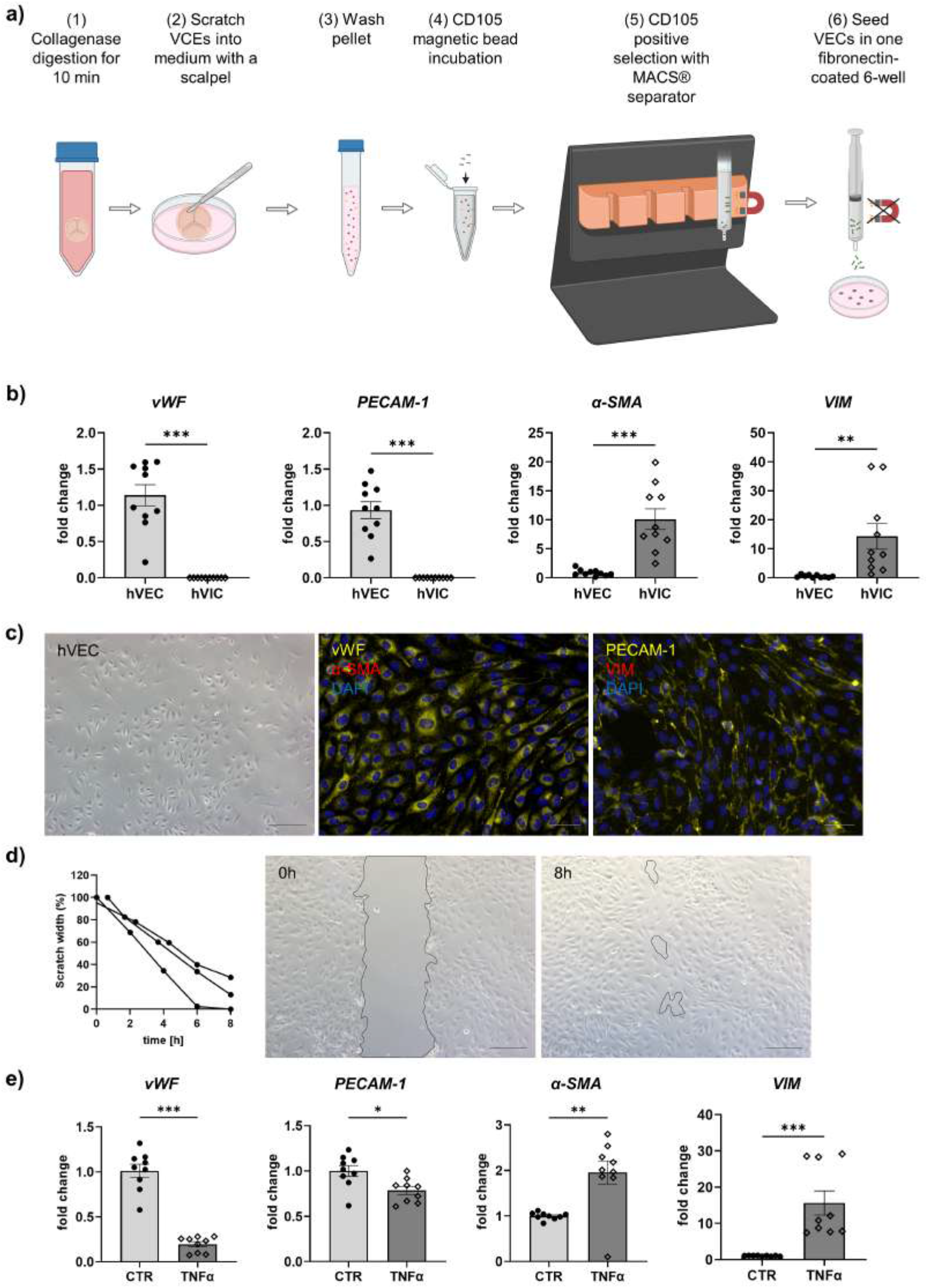
Human valvular endothelial cell isolation and characterization. **(a)** Workflow of human VEC isolation: (1) The explanted aortic valve cusps were incubated for 10 minutes in collagenase Type II solution. (2) VECs were carefully scratched with a scalpel into a dish filled with VEC medium, which (3) was transferred to a 15 ml tube and washed twice for 15 minutes at 1000 rpm at room temperature. (4) Cells were incubated with CD105 magnetic beads for 15 minutes and (5) the CD105 positive selection was carried out with MACS^®^ separator, according to the manufacturer’s protocol. (6) The pure VEC population was seeded in one well of a fibronectin-coated 6-well plate. **(b)** Gene expression analysis of isolated human valvular endothelial cells (hVECs) displayed elevated endothelial marker expression of von Willebrand factor (*vWF*) and platelet endothelial adhesion molecule-1 (*PECAM-1*) and a lower expression of the mesenchymal markers alpha smooth muscle actin (*α-SMA*) and vimentin (*VIM*) compared to human valvular interstitial cells (hVICs). **(c)** Brightfield and immunofluorescence images of VECs positive for vWF and PECAM-1 and negative for α-SMA and VIM. **(d)** Migration analysis by scratch wound assay showed a scratch closure from 100% to 15% after 8 hours. **(e)** Inducing EndMT by incubating VECs with tumor necrosis factor alpha (TNFα lead to an upregulation of α-SMA and VIM, and a downregulation of PECAM-1 and vWF. (b) n=5 donors. (c-e) n=3 donors, *P < 0.05, **P < 0.01, ***P < 0.001, ns not-significant, analyzed by Student t-test, 2 tail, unpaired, scale bar 200μm, (a) created with BioRender.com.

MACS-sorted hVICs were characterized by RT-qPCR to quantify interstitial marker gene expression (Figure 2a). Whereas *α-SMA* and *VIM* expression were significantly upregulated, *vWF* as well as *PECAM-1* showed decreased expression values compared to hVECs (Figure 1b). In immunofluorescence staining of hVICs a positive staining for intracellularly distributed α-SMA and VIM was observed and expression of vWF and PECAM-1 was not detected (Figure 2b). To analyze the migration properties of hVICs, a scratch wound was performed, and the closure observed for 8 hours. The remaining scratch area was 87% after 8 hours (Figure 2c). *In vitro* calcification, by incubating hVICs with OM or PCM for 7 days revealed an overexpression of alkaline phosphatase (*ALPL*), a hallmark of calcification, in OM but not in PCM (Figure 2d). In addition, other calcification-related genes, for instance, osteocalcin (*BGLAP*) and runt-related factor 2 (*RUNX2*), were upregulated in both conditions. Alizarin red staining of calcified cells provided a further layer of confirmation that the isolated cells were VICs with osteoblastic differentiation capacity (figure 2e). These data also confirmed that *in vitro* calcification of isolated hVICs could be induced successfully.

**Figure 2:**
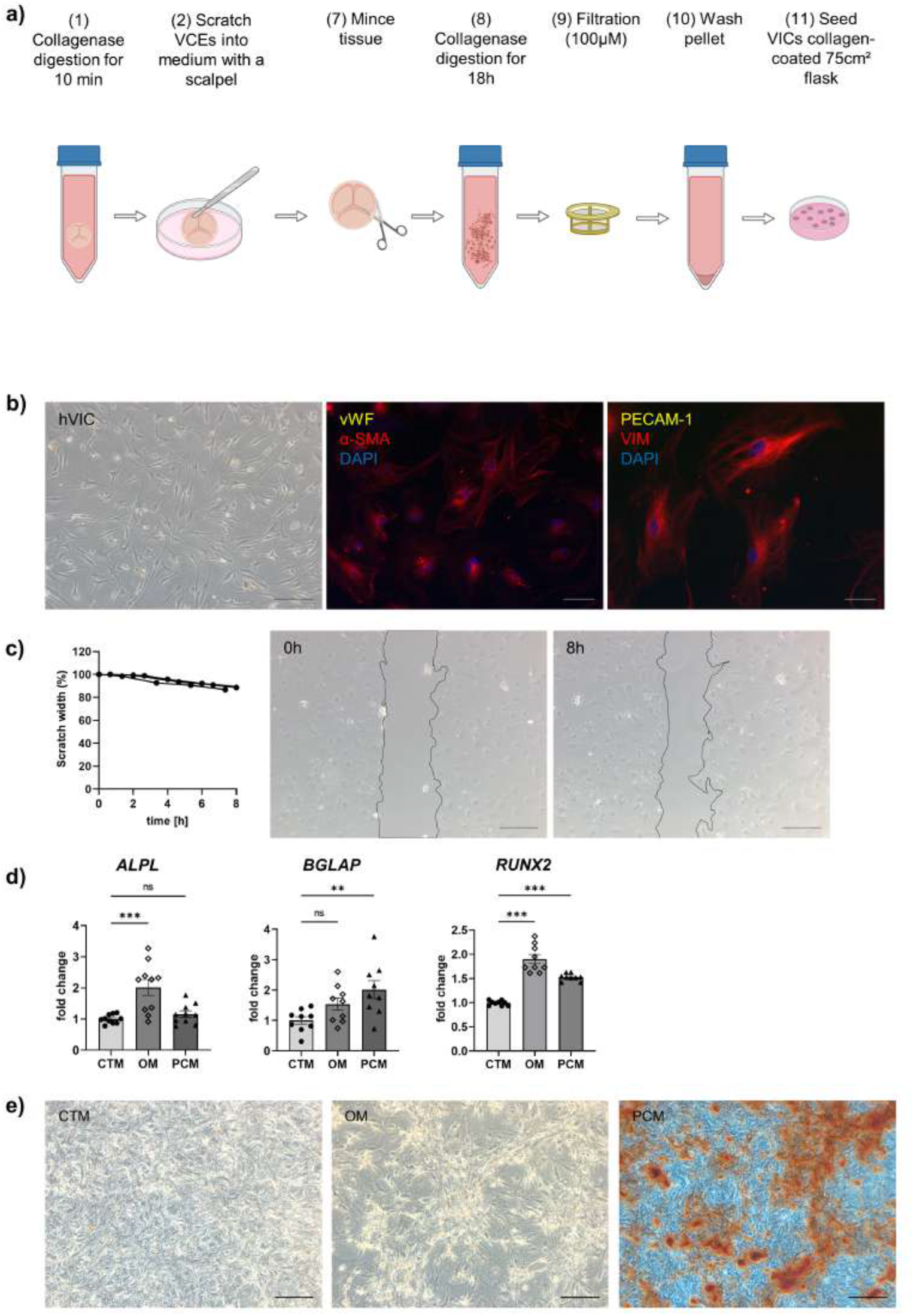
Human valvular interstitial cells isolation and characterization. **(a)** Workflow of human VIC isolation: (7) the remaining aortic valve tissue after VEC removal was minced into small pieces and (8) further digested in collagenase solution for 18 hours. (9) Cell suspension was filtered (100μM nylon mesh filter) and (10) the pellet washed twice. (11) VICs were seeded in a collagen-coated 75cm² flask. **(b)** Brightfield images and immunofluorescence staining of hVICs positive for mesenchymal markers alpha smooth muscle actin (*α-SMA*) and vimentin (*VIM*) and negative for von Willebrand factor (*vWF*) and platelet endothelial adhesion molecule-1 (*PECAM-1*). **(c)** Migration analysis by scratch wound assay showed a scratch closure from 100% to 15% in 8h. **(d)** Gene expression analysis of hVICs incubated with osteogenic medium (OM) or pro-calcifying medium (PCM) for 7 days to induced *in vitro* calcification. QPCR analysis showed increased alkaline phosphatase (*ALPL*) expression in OM, but not in PCM. Osteocalcin (*BGLAP*) and bone morphogenic protein 2 (*BMP2*) expression was upregulated in OM and PCM, compared to control medium (CTM). **(e)** Alizarin red staining displayed increased calcium nodule formation in OM and PCM. n=3 donors, *P < 0.05, **P < 0.01, ***P < 0.001, ns not-significant, (b, c) analyzed by Student t-test, 2 tail, unpaired, (d, e) analyzed by one-way ANOVA with Bonferroni’s correction, scale bar 200μm, (a) created with BioRender.com.

### Porcine aortic valve cell isolation revealed specific VEC and VIC cell populations

To assess whether porcine cells can be used alternatively for cell culture experiments instead of difficult-to-obtain human cells, porcine VECs (pVECs) and porcine VICs (pVICs) were characterized. Cells were isolated using the same protocol as for human cells (Figure 3a). Analysis of characteristic marker gene expression in isolated pVECs revealed elevated *vWF* and *PECAM-1* levels and a reduction of *α-SMA* compared to pVICs. *VIM* was also upregulated in pVECs (Figure 3b). In immunofluorescence staining, positive staining for vWF was observed but no α-SMA signal was detectable in pVECs (Figure 3c). To assess the migration behavior of hVECs, a scratch wound assay was performed which revealed a scratch closure to 70% after 8h (Figure 3d). This data provided evidence that pVECs displayed an endothelial specific phenotype. However, *in vitro* EndMT induction using TNFα demonstrated a decrease of EndMT markers (Figure 3e), indicating that cells had not undergone efficient transdifferentiation.

**Figure 3:**
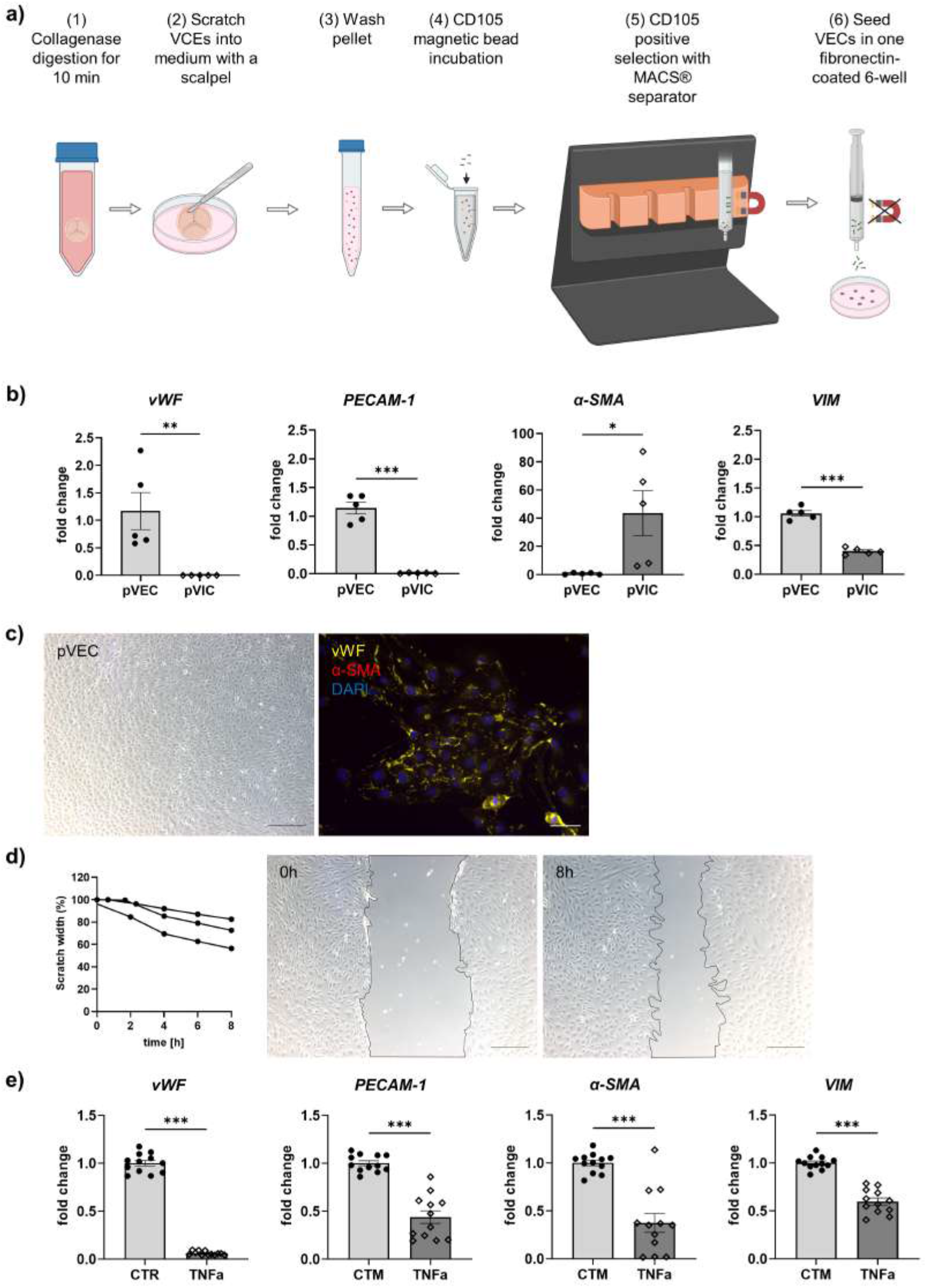
Isolation and characterization of porcine valvular endothelial cells. **(a)** Workflow of porcine VEC isolation. **(b)** Gene expression analysis of pVECs revealed a significant upregulation of v*o*n Willebrand factor (*vWF*), *PECAM-1* and vimentin (*VIM*) and a downregulation of alpha smooth muscle actin (*α-SMA*), compared to pVICs. **(c)** Brightfield images and immunofluorescence staining of pVECs showing a positive signal for vWF and an absent signal for α-SMA. **(d)** Analysis of migration properties showed a scratch closure to 70% after 8 hours. **(e)** In vitro EndMT induction by TNFα lead to a downregulation of endothelial (*PECAM-1, vWF*) and mesenchymal markers (*α-SMA, VIM*). (b) n=5 donors. (c-e) n=3 donors, *P < 0.05, **P < 0.01, ***P < 0.001, ns not-significant, analyzed by Student t-test, 2 tail, unpaired, scale bar 200μm, (a) created with BioRender.com.

To characterize isolated pVICs, using the same protocol as for human cells (Figure 4a), characteristic interstitial cell markers were analyzed, showing an upregulation of *α-SMA* in pVICs compared to pVECs (Figure 3b). Noteworthily, endothelial markers, *vWF* and *PECAM-1* as well as *VIM* were decreased. Immunofluorescence staining displayed a positive signal for α-SMA and VIM, while vWF and PECAM-1 were undetectable (Figure 4b). Scratch-wound based functional assays revealed a scratch closure after 8 hours of 87% (Figure 5c). These data suggest a VIC specific phenotype of the isolated cells. To assess whether calcification can be induced in pVICs *in vitro*, cells were incubated with OM or PCM for 7 days, showing a significant upregulation of calcification-related genes, namely, *ALPL, BMP2, BGLAP* and osteopontin *(SPP1*) in OM and PCM (Figure 5d). Additionally, alizarin red staining revealed enhanced calcific nodule staining in OM treated and more pronounced staining in PCM treated cells, due to deposition of calcium phosphate (Ca_3_(PO_4_)_2_) into the cellular matrix (Figure 5e). This data provided evidence that pVICs can be utilized for *in vitro* calcification experiments.

**Figure 4:**
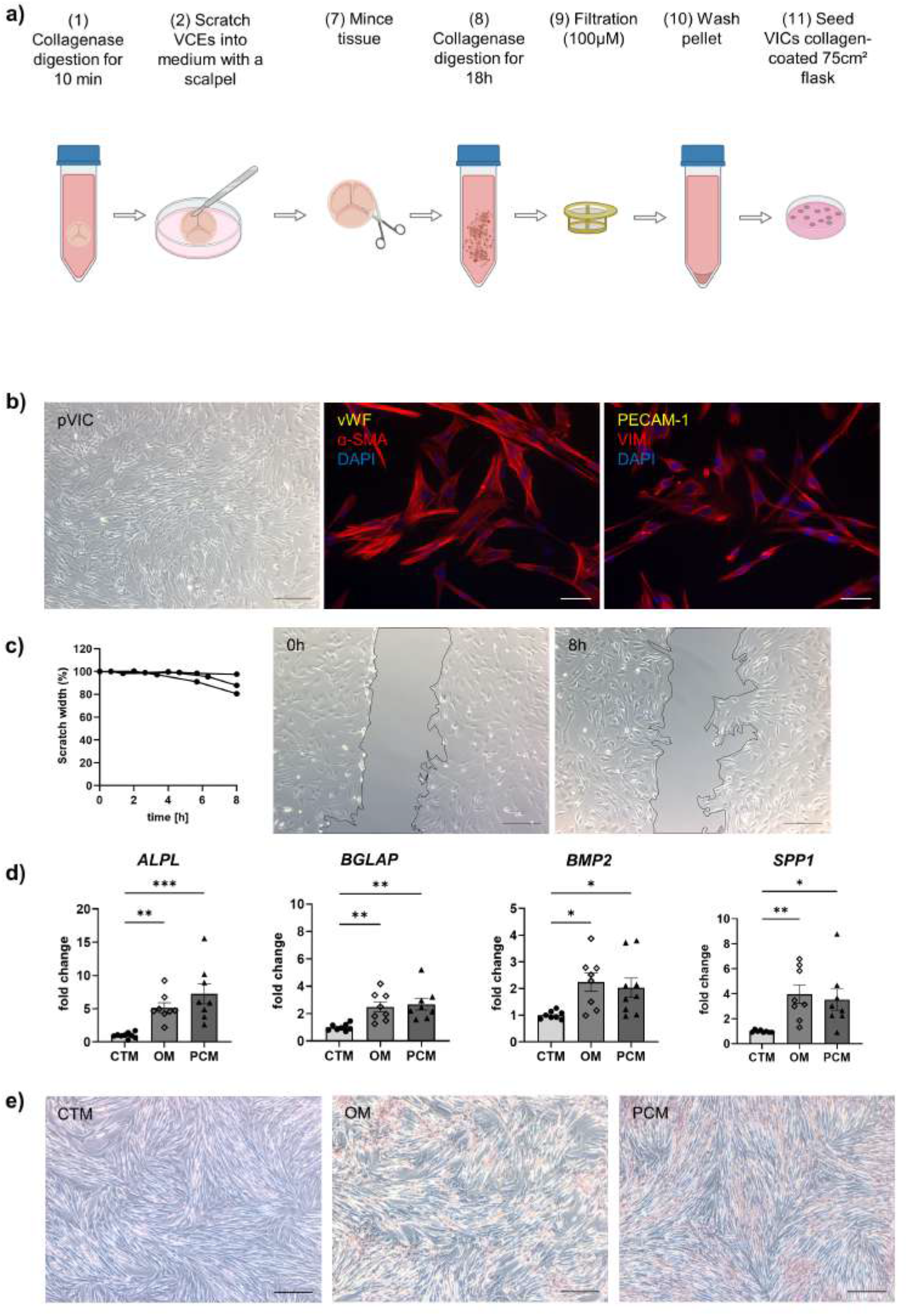
Isolation and characterization of porcine interstitial cells. **(a)** Workflow of porcine VIC isolation. **(b)** Brightfield and immunofluorescence images of pVICs displaying a positive staining for a-SMA and VIM and a negative signal for PECAM-1 and vWF. **(c)** Migration assays revealed a scratch closure to 87% after 8 hours. **(d)** Gene expression analysis of pVICs were treated with osteogenic medium (OM) or pro-calcifying medium (PCM) for 7 days to induce *in vitro* calcification. Calcific markers, such as alkaline phosphatase (*ALPL*), bone morphogenic protein 2 (*BMP2*), osteocalcin (*BGLAP*) and osteopontin (*SPP1*) were significantly upregulated in OM and PCM. **(e)** Alizarin Red staining showed pronounced calcific nodules in OM and PCM, compared to control medium (CTM). n=3 donors, *P < 0.05, **P < 0.01, ***P < 0.001, ns not-significant, (b, c) analyzed by Student t-test, 2 tail, unpaired, (d) analyzed by one-way ANOVA with Bonferroni’s correction, scale bar 200μm, (a) created with BioRender.com.

**Figure 5:**
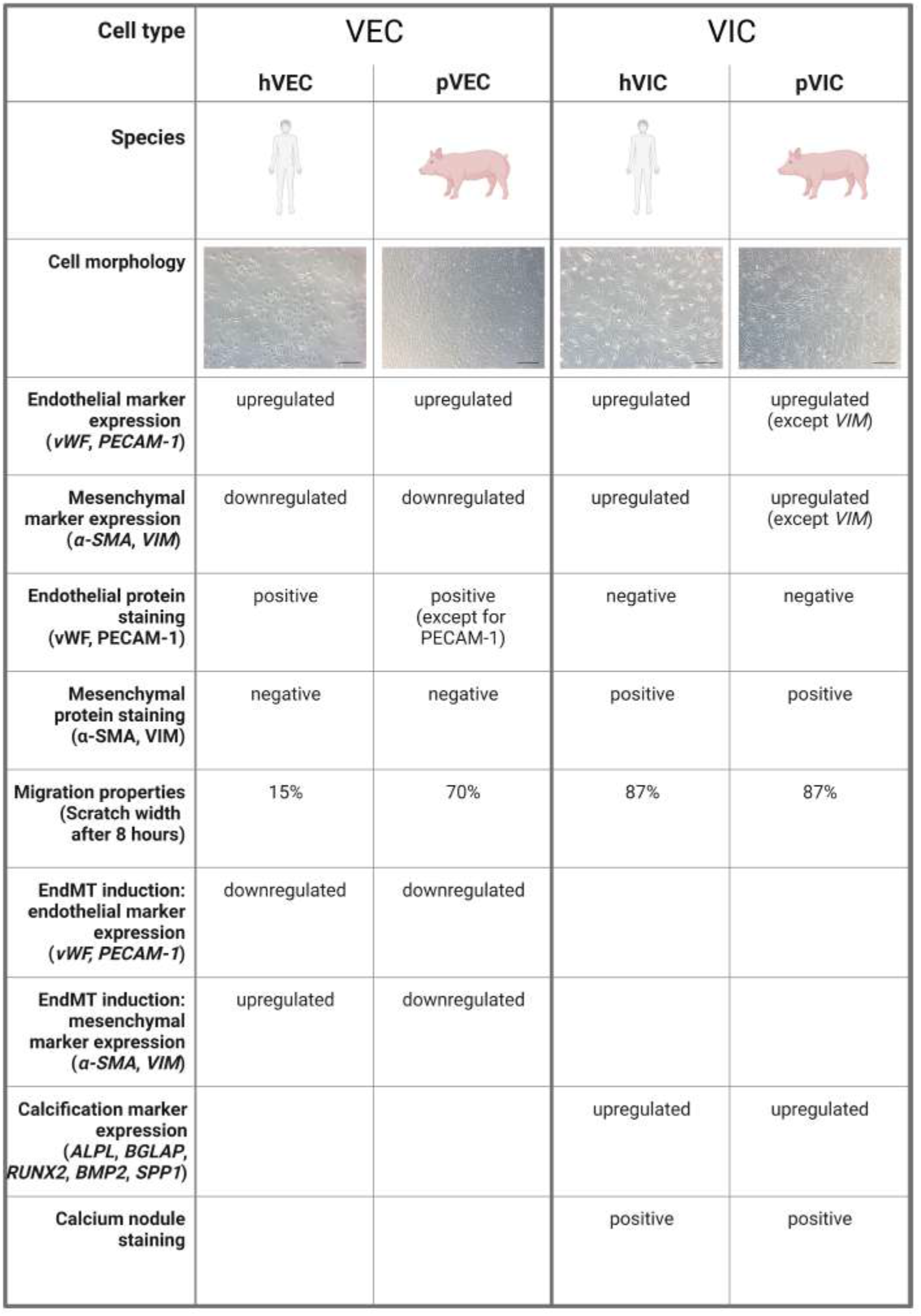
A comparison of human and porcine valvular endothelial and interstitial cells. Valvular endothelial cells (VECs), valvular interstitial cells (VICs), human VECs (hVECs), human VICs (hVICs), porcine VECs (pVECs), porcine VICs (pVICs), von Willebrand factor (vWF), platelet adhesion molecule-1 (PECAM-1), alpha smooth muscle actin (α-SMA), vimentin (VIM), alkaline phosphatase (ALPL), osteocalcin (BGLAP), runt-related factor X 2 (RUNX2), bone morphogenic protein 2 (BMP2), osteopontin (SPP1).

## Discussion

Analyzing aortic valve cells is of major relevance for a better understanding of the pathology of aortic valve stenosis, but obtaining, isolating and cultivating these cells is a demanding task. Human aortic valve tissue is difficult to obtain, and the acquisition of non-stenotic control tissues is often especially challenging. In principle, living aortic valve cells can be gained from donor hearts that are unsuitable for transplantation, as long as the aortic valve is not affected by underlying disease [29]. This is the most suitable source of control tissue, but since there are only few transplant units and a limited number of donors who would be appropriate candidates, this material is difficult to obtain [30, 31]. It is also possible to use valves that were explanted because of aortic dissection, since this is presumed to not affect the aortic valve [32]. However, these operations are typically performed as emergency operations, with ensuing problems concerning both informed consent and the isolation procedure. The most common source of non-calcified human aortic valve tissue are patients undergoing SAVR due to aortic regurgitation in the context of aortic root or ascending aortic aneurysm.

Standardizing the method of aortic valve cell isolation is crucial to ensure optimal data exchange between investigators. In a first attempt to standardize cell isolation from human and porcine aortic valve tissues, we propose a method based on previous publications with some refinements [17, 19, 23, 33, 34]. We removed VECs from AVS cusps via scraping cells from valve tissue by the careful use of a scalpel prior to collecting and affinity-based isolation. This increased the number of VECs gained from the isolation process, when compared to removal with a cotton swab. Moreover, we added a magnetic-activated cell sorting (MACS, Miltenyi Biotech) step with the specific endothelial marker endoglin (CD105) in order to ensure a pure VEC isolation [22]. This endothelial cell separation has previously been described for human vascular endothelial cells. Based on our study, plating VICs on a collagen-coated 75 cm^2^ culture flask is more appropriate for cell culture-based experiments [17, 20]. Due to the thorough sorting and lesser number of VECs in general, VEC yields are significantly lower than VIC yields and thus, porcine or human cells can be plated on one fibronectin-coated 6-well to enable cell-cell interaction [17].

The second aim of this manuscript was to compare human and porcine VECs and VICs with each other to assess whether easier-to-obtain porcine aortic valve tissues and their cells can be used for primary *in vitro* investigations to study human AVS pathologies. We found commensurabilities but also disparities between human and porcine aortic valve cells. Under physiological conditions hVECs and pVECs resemble endothelial cells covering the vasculature displaying a typical cuboid, endothelial cell shape with cell-junctions and expression of specific endothelial markers such as *PECAM-1* and *vWF* [35]. To test whether the differentiation of VECs to mesenchymal cells by TNFα treatment can be performed successfully, gene expression analysis of EndMT markers, such as *α-SMA* and *VIM* were performed after 7 days of TNFα treatment [36]. In hVECs incubated with TNFα, the expression of mesenchymal cell markers such as *α-SMA* and *VIM* was increased, whereas *vWF* and *PECAM-1* were decreased. In pVECs the endothelial makers were also downregulated but surprisingly, this also applies for *α-SMA* and *VIM*. During this transition, activated VECs progressively lose their endothelial properties, including cell junctions, and their specific endothelial markers, display increased cellular invasion and acquire a mesenchymal phenotype with improved migratory capabilities[35]. hVICs and pVICs displayed the typical spindle-shaped VIC morphology with a swirling pattern that start to grow over each other at confluency [23]. VICs are a heterogenous and highly plastic cell population residing within the tissue matrix of the aortic valve [37, 38]. In healthy human aortic valves, most VICs are considered to be quiescent with a fibroblast-like phenotype, which can be distinguished from VECs by positive expression of *VIM* and the absence of endothelial markers *vWF* and *PECAM-1* [39–41]. Upon activation they can undergo differentiation into activated myofibroblast-like VICs, which are, besides vimentin, also positive for α-SMA [37, 42]. This leads to the conclusion that hVICs display a myofibroblastic-like phenotype. Intriguingly, comprehensive studies identified cellular heterogeneities in aortic valve tissues and identified 234 cell clusters, representing 58 major cell types [43]. This study also reported endothelial, and interstitial cell heterogeneities, including endothelial phenotype during EndMT via in-depth integrated multi-omics, [43]. Single-cell RNA sequencing and proteomic analysis from pVECs undergoing EndMT could provide further insight into the potential genetic differences. To this end, our findings are also in line with above-mentioned report, suggesting a higher degree of cellular heterogeneity in the isolated porcine cell population.

Although the isolation, cultivation, and *in vitro* experiments with aortic valve cells are challenging, many advances have been made in recent years [17-18]. Different methods for isolation, EndMT, and calcification have been established around the world [18]. However, there is currently no reproducible experimental standardization, which would be crucial to ensure optimal comparisons between laboratories and to connect all findings. This manuscript aims to initiate the first step towards standardization of human and porcine aortic valve cell isolation. *In vitro* experiments with the aortic valve, cells are immensely important to improve our understanding of involved pathomechanisms during the development of CAVD and will undoubtedly lead to the identification of druggable targets. Additionally, no direct and detailed comparison between human and porcine aortic valve tissue has been reported.

However, our study has some limitations, namely, the patient samples were all collected at the University of Bonn, Germany, thus, the patient-derived data presented is limited to the Caucasian race. Therefore, no information on race-dependent differences and heterogeneities were derivable in isolated cells from AV tissues of AVS patients who underwent SAVR. Although the characteristic gene expression levels of VIC and VEC markers are similar in patient-derived valvular cells there are some differences in cells of porcine origin, as confirmed by RT-qPCR. Due to the technological limitations of valve cell isolations, it is still very challenging to completely confirm that valvular cell populations are absolutely pure and without any other contamination in the isolates. In cardiovascular research, a wide range of studies has reported dynamic alterations in the expression of genes during different cardiovascular conditions, leading to an interest in single-cell resolution investigations[44-46]. Regarding patients with AVS, only some patients display impaired heart function, as evidenced by a reduction in LVEF (left ventricular ejection fraction, indicating the severity of the pathological changes of the failing heart, while other patients remain without LVEF impairment, demonstrating heterogeneity of pathological changes during AVS[47-48]. To enumerate whether human and porcine cells are comparable in terms of cellular and other biological properties of valvular cells, we summarized our findings in figure 5. Such limitations can be accounted for in the case of individual experiments in specific lab settings in cardiovascular research.

In conclusion, our findings offer new avenues into alternative cellular model systems that can be used in a wide range of biomedical research. We demonstrated for the first time in detail, that 1) patient-derived AV cells, namely, VICs and VECs, isolated from AV tissue explants from AVS patients can be isolated as a pure population. These cells express typical endothelial- and interstitial cell markers and are comparable morphologically; 2) patient-derived cells can be maintained in culture for a prolonged time (passages, 5-6), and possess the potential for usage in downstream experiments; 3) functional analysis demonstrated that isolated cells retain endothelial- and interstitial cell properties/function (for instance in VEC: migration, tube formation, EndMT induction; in VIC: *in vitro* calcification to facilitate osteoblastic differentiation upon induction with OM or PCM) 4) a comparative and reproducible analysis of human vs. porcine cells revealed that both cell types are comparable, although some cellular heterogeneity in VECs of porcine origin (pVECs) exist.

In summary, our study has revealed an easy-to-obtain and financially viable alternative cellular model system that can be used in cardiovascular research. Our findings can also help circumvent limitations of accessibility and challenges in obtaining human AV tissues and provide a viable solution for lab-usable cellular models in aortic valve research.

## Acknowledgements

The authors would like to thank Anna Flender for technical support and the heart surgeons at the Heart Center of University Hospital Bonn for providing the aortic valve tissue. The graphical images were created with *BioRender*.*com*.

## Funding

DN, FJ and GN are supported by the DFG, SFB TRR259/1 [Sonderforschungsbereich/Transregio] (397484323). PRG is funded by the Else-Kröner-Fresenius Foundation. KM, MRH, and FJ are funded by the Corona foundation and German Society of Cardiology (DGK). EA is supported by National Institute of Health (NIH) grants, R01HL136431, R01HL147095, and R01HL141917.

## Figures with legends

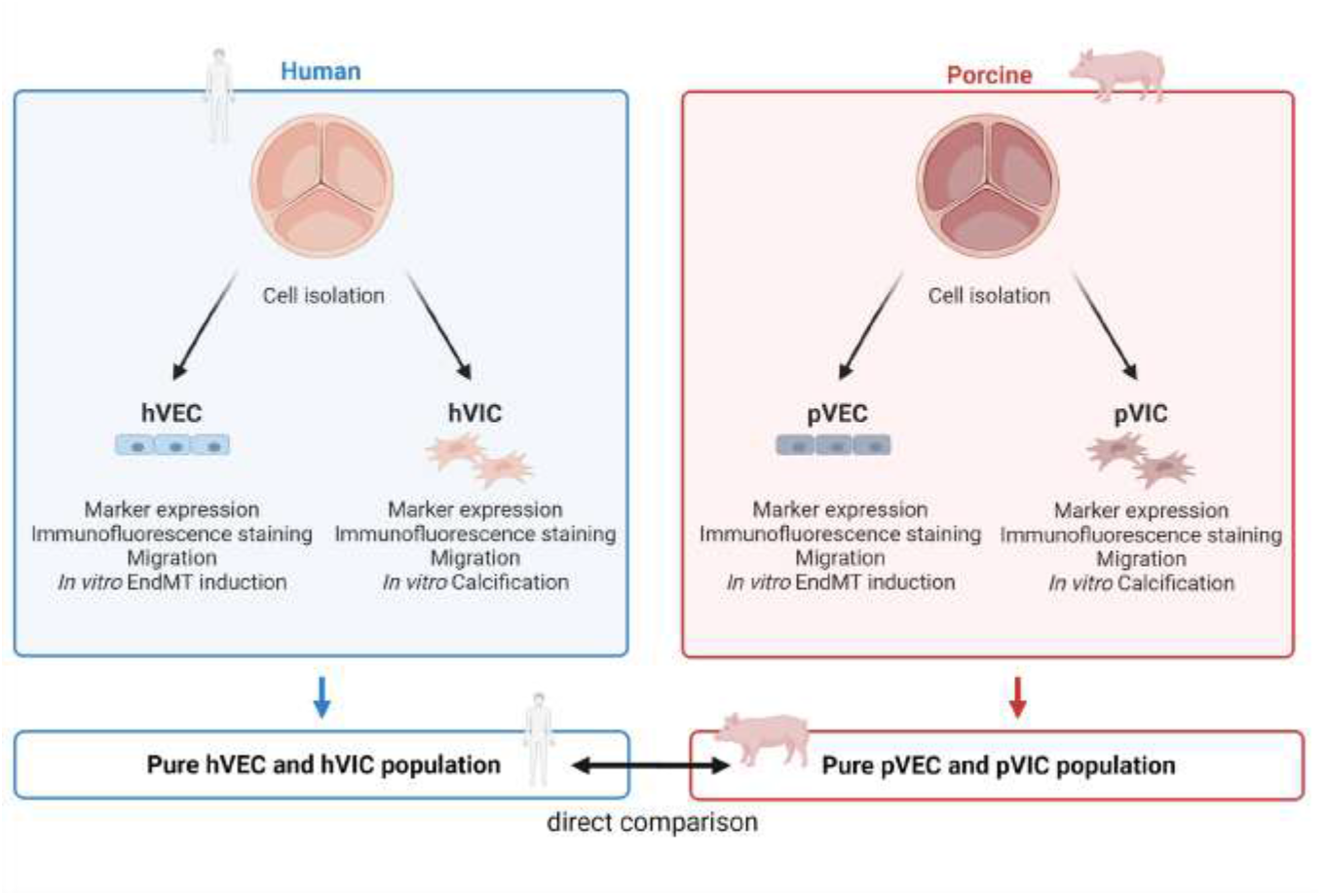

**Graphical abstract: *Aortic valve cell isolation: a comparative study of endothelial and interstitial cell isolation, and characterization with valvular tissue from porcine and humans***.

Human aortic valve endothelial cells (hVECs) and interstitial cells (hVICs) as well as porcine aortic valve endothelial cells (pVECs) and interstitial cells (pVICs) were isolated from human explants from patients undergoing surgical aortic valve replacement or porcine valvular tissue, respectively. To ensure specific cell populations, cells were characterized by marker expression analysis, immunofluorescence staining as well as migration assays. Additionally, endothelial-to-mesenchymal transition (EndMT) was induced *in vitro* in hVECs and pVECs along with *in vitro* calcification of hVICs and pVICs. Finally, these results were compared from human cells to pig cells. To date, no direct comparison has been performed.

## Notes

### Competing Interest Statement

The authors have declared no competing interest.

## References

1. Osnabrugge RLJ, Mylotte D, Head SJ, van Mieghem NM, Nkomo VT, LeReun CM, Bogers AJJC, Piazza N, Kappetein AP (2013) Aortic stenosis in the elderly: disease prevalence and number of candidates for transcatheter aortic valve replacement: a meta-analysis and modeling study. Journal of the American College of Cardiology 62(11):1002–1012

2. Kodali SK, Williams MR, Smith CR, Svensson LG, Webb JG, Makkar RR, Fontana GP, Dewey TM, Thourani VH, Pichard AD, Fischbein M, Szeto WY, Lim S, Greason KL, Teirstein PS, Malaisrie SC, Douglas PS, Hahn RT, Whisenant B, Zajarias A, Wang D, Akin JJ, Anderson WN, Leon MB (2012) Two-Year Outcomes after Transcatheter or Surgical Aortic-Valve Replacement. The New England Journal of Medicine 366(18):1686–1695

3. Nkomo VT, Gardin JM, Skelton TN, Gottdiener JS, Scott CG, Enriquez-Sarano M (2006) Burden of valvular heart diseases: a population-based study. The Lancet 368(9540):1005–1011

4. Mönckeberg JG (1904) Der normale histologische Bau und die Sklerose der Aortenklappen. Virchows Archiv für pathologische Anatomie und Physiologie und für klinische Medizin 176):472–514

5. Selzer A (1987) Changing Aspects of the Natural History of Valvular Aortic Stenosis. New England Journal of Medicine 317):91–98

6. Aikawa E, Nahrendorf M, Figueiredo J-L, Swirski FK, Shtatland T, Kohler RH, Jaffer FA, Aikawa M, Weissleder R (2007) Osteogenesis associates with inflammation in early-stage atherosclerosis evaluated by molecular imaging in vivo. Circulation 116(24):2841–2850

7. Rajamannan NM, Evans FJ, Aikawa E, Grande-Allen KJ, Demer LL, Heistad DD, Simmons CA, Masters KS, Mathieu P, O’Brien KD, Schoen FJ, Towler DA, Yoganathan AP, Otto CM (2011) Calcific aortic valve disease: not simply a degenerative process: A review and agenda for research from the National Heart and Lung and Blood Institute Aortic Stenosis Working Group. Executive summary: Calcific aortic valve disease-2011 update. Circulation 124(16):1783–1791

8. Makkar Raj R., Fontana Gregory P., Jilaihawi Hasan, Kapadia Samir, Pichard Augusto D., Douglas Pamela S., Thourani Vinod H., Babaliaros Vasilis C., Webb John G., Herrmann Howard C., Bavaria Joseph E., Kodali Susheel, Brown David L., Bowers Bruce, Dewey Todd M., Svensson Lars G., Tuzcu Murat, Moses Jeffrey W., Williams Matthew R., Siegel Robert J., Akin Jodi J., Anderson William N., Pocock Stuart, Smith Craig R., Leon Martin B. (2012) Transcatheter Aortic-Valve Replacement for Inoperable Severe Aortic Stenosis. New England Journal of Medicine 366(18):1696–1704

9. Otto CM, Kuusisto J, Reichenbach DD, Gown AM, O’Brien KD (1994) Characterization of the early lesion of ‘degenerative’ valvular aortic stenosis. Histological and immunohistochemical studies. Circulation 90):844–853

10. Yearwood TL, Misbach GA, Chandran KB (1989) Experimental fluid dynamics of aortic stenosis in a model of the human aorta. Clinical Physicy and Physiological Measurement 10(11)

11. New SEP, Aikawa E (2011) Molecular imaging insights into early inflammatory stages of arterial and aortic valve calcification. Circulation research 108(11):1381–1391

12. Timmerman LA, Grego-Bessa J, Raya A, Bertrán E, Pérez-Pomares JM, Díez J, Aranda S, Palomo S, McCormick F, Izpisúa-Belmonte JC, La Pompa JL de (2004) Notch promotes epithelial-mesenchymal transition during cardiac development and oncogenic transformation. Genes & development 18(1):99–115

13. Wirrig EE, Yutzey KE (2014) Conserved transcriptional regulatory mechanisms in aortic valve development and disease. Arteriosclerosis, thrombosis, and vascular biology 34(4):737–741

14. Kovacic JC, Dimmeler S, Harvey RP, Finkel T, Aikawa E, Krenning G, Baker AH (2019) Endothelial to Mesenchymal Transition in Cardiovascular Disease: JACC State-of-the-Art Review. Journal of the American College of Cardiology 73(2):190–209

15. Olsson M, Dalsgaard C-J, Haegerstrand A, Rosenqvist M, Rydén L, Nilsson J (1994) Accumulation of T lymphocytes and expression of interleukin-2 receptors in nonrheumatic stenotic aortic valves. Histological and immunohistochemical studies. Journal of the American College of Cardiology 23(5):1162–1170

16. Goody PR, Hosen MR, Christmann D, Niepmann ST, Zietzer A, Adam M, Bönner F, Zimmer S, Nickenig G, Jansen F (2020) Aortic Valve Stenosis: From Basic Mechanisms to Novel Therapeutic Targets. Arteriosclerosis, thrombosis, and vascular biology 40(4):885–900

17. Gould RA, Butcher JT (2010) Isolation of valvular endothelial cells. Journal of visualized experiments : JoVE 46)

18. Johnson CM, Fass DN (1983) Porcine cardiac valvular endothelial cells in culture. A relative deficiency of fibronectin synthesis in vitro. Laboratory investigation; a journal of technical methods and pathology 49(5):589–598

19. El Husseini D, Boulanger M-C, Mahmut A, Bouchareb R, Laflamme M-H, Fournier D, Pibarot P, Bossé Y, Mathieu P (2014) P2Y2 receptor represses IL-6 expression by valve interstitial cells through Akt: implication for calcific aortic valve disease. Journal of molecular and cellular cardiology 72):146–156

20. Goto S, Rogers MA, Blaser MC, Higashi H, Lee LH, Schlotter F, Body SC, Aikawa M, Singh SA, Aikawa E (2019) Standardization of Human Calcific Aortic Valve Disease in vitro Modeling Reveals Passage-Dependent Calcification. Frontiers in cardiovascular medicine 6):49

21. O’Brien KD (2007) Epidemiology and genetics of calcific aortic valve disease. Journal of investigative medicine : the official publication of the American Federation for Clinical Research 55(6):284–291

22. Wang H, Sridhar B, Leinwand LA, Anseth KS (2013) Characterization of cell subpopulations expressing progenitor cell markers in porcine cardiac valves. PloS one 8(7):e69667

23. Johnson CM, Hanson MN, Helgeson SC (1987) Porcine cardiac valvular subendothelial cells in culture: Cell isolation and growth characteristics. J Mol Cell Cardiol 19):1158–1193

24. Naimark WA, Lee JM, Limeback H, Cheung DT (1992) Correlation of structure and viscoelastic properties in the pericardia of four mammalian species. Heart and Circulatory Physiology 263(4)

25. Merryman WD, Youn I, Lukoff HD, Krueger PM, Guilak F, Hopkins RA, Sacks MS (2006) Correlation between heart valve interstitial cell stiffness and transvalvular pressure: implications for collagen biosynthesis. American journal of physiology. Heart and circulatory physiology 290(1):H224–31

26. Rutkovskiy A, Malashicheva A, Sullivan G, Bogdanova M, Kostareva A, Stensløkken K-O, Fiane A, Vaage J (2017) Valve Interstitial Cells: The Key to Understanding the Pathophysiology of Heart Valve Calcification. Journal of the American Heart Association 6(9)

27. World Medical Association (2001) Declaration of Helsinki. World Medical Association Declaration of Helsinki. Ethical principles for medical research involving human subjects. Bulletin of the World Health Organization 79(4):373–374

28. Liang C-C, Park AY, Guan J-L (2007) In vitro scratch assay: a convenient and inexpensive method for analysis of cell migration in vitro. Nature protocols 2(2):329–333

29. Messer SJ, Axell RG, Colah S, White PA, Ryan M, Page AA, Parizkova B, Valchanov K, White CW, Freed DH, Ashley E, Dunning J, Goddard M, Parameshwar J, Watson CJ, Krieg T, Ali A, Tsui S, Large SR (2016) Functional assessment and transplantation of the donor heart after circulatory death. The Journal of heart and lung transplantation : the official publication of the International Society for Heart Transplantation 35(12):1443–1452

30. Ferdous Z, Jo H, Nerem RM (2011) Differences in valvular and vascular cell responses to strain in osteogenic media. Biomaterials 32(11):2885–2893

31. Yang X, Fullerton DA, Su X, Ao L, Cleveland JC, Meng X (2009) Pro-osteogenic phenotype of human aortic valve interstitial cells is associated with higher levels of Toll-like receptors 2 and 4 and enhanced expression of bone morphogenetic protein 2. Journal of the American College of Cardiology 53(6):491–500

32. Wang B, Li F, Zhang C, Wei G, Liao P, Dong N (2016) High-mobility group box-1 protein induces osteogenic phenotype changes in aortic valve interstitial cells. The Journal of thoracic and cardiovascular surgery 151(1):255–262

33. Rush MN, Coombs KE, Hedberg-Dirk EL (2015) Surface chemistry regulates valvular interstitial cell differentiation in vitro. Acta biomaterialia 28):76–85

34. Cheung W-Y, Young EW, Simmons CA (2008) Techniques for Isolating and Purifying Porcine AorticValve Endothelial Cells. The Journal of Heart Valve Disease 17):647–681

35. Ma X, Zhao D, Yuan P, Li J, Yun Y, Cui Y, Zhang T, Ma J, Sun L, Ma H, Zhang Y, Zhang, Haiuhou, Zhang W, Huang J, Zou C, Wang Z (2020) Endothelial-to-Mesenchymal Transition in Calcific Aortic Valve Disease. Acta Cardiol Sin 23(3)

36. Dejana E, Hirschi KK, Simons M (2017) The molecular basis of endothelial cell plasticity. Nature communications 8):14361

37. Hjortnaes J, Shapero K, Goettsch C, Hutcheson JD, Keegan J, Kluin J, Mayer JE, Bischoff J, Aikawa E (2015) Valvular interstitial cells suppress calcification of valvular endothelial cells. Atherosclerosis 242(1):251–260

38. Liu AC, Joag VR, Gotlieb AI (2007) The emerging role of valve interstitial cell phenotypes in regulating heart valve pathobiology. The American Journal of Pathology 171(5):1407–1418

39. Messier RH, Bass BL, Aly HM, Jones JL, Domkowski PW, Wallace RB, Hopkins RA (1994) Dual structural and functional phenotypes of the porcine aortic valve interstitial population: characteristics of the leaflet myofibroblast. Journal of surgical research 57):1–21

40. Farrar EJ, Butcher JT (2014) Heterogeneous susceptibility of valve endothelial cells to mesenchymal transformation in response to TNFα. Annals of biomedical engineering 42(1):149–161

41. Porras AM, van Engeland NCA, Marchbanks E, McCormack A, Bouten CVC, Yacoub MH, Latif N, Masters KS (2017) Robust Generation of Quiescent Porcine Valvular Interstitial Cell Cultures. Journal of the American Heart Association 6(3)

42. Rabkin E, Aikawa M, Stone JR, Fukumoto Y, Libby P, Schoen FJ (2001) Activated interstitial myofibroblasts express catabolic enzymes and mediate matrix remodeling in myxomatous heart valves. Circulation 104(21):2525–2532

43. Wang, F., Ding, P., Liang, X., Ding, X., Brandt, C.B., Sjöstedt, E., Zhu, J., Bolund, S., Zhang, L., de Rooij, L.P. and Luo, L., 2022. Endothelial cell heterogeneity and microglia regulons revealed by a pig cell landscape at single-cell level. Nature communications, 13(1), pp.1–18.

44. Hulsmans, M., Sinnaeve, P., Van der Schueren, B., Mathieu, C., Janssens, S. and Holvoet, P., 2012. Decreased miR-181a expression in monocytes of obese patients is associated with the occurrence of metabolic syndrome and coronary artery disease. The Journal of Clinical Endocrinology & Metabolism, 97(7), pp.E1213–E1218

45. Nightingale, A.K. and Horowitz, J.D., 2005. Aortic sclerosis: not an innocent murmur but a marker of increased cardiovascular risk. Heart, 91(11), pp.1389–1393.

46. Otto, C.M. and Prendergast, B., 2014. Aortic-valve stenosis—from patients at risk to severe valve obstruction. New England Journal of Medicine, 371(8), pp.744-756.

47. Makkar, R.R., Fontana, G.P., Jilaihawi, H., Kapadia, S., Pichard, A.D., Douglas, P.S., Thourani, V.H., Babaliaros, V.C., Webb, J.G., Herrmann, H.C. and Bavaria, J.E., 2012. Transcatheter aortic-valve replacem

48. Wirrig, E.E. and Yutzey, K.E., 2014. Conserved transcriptional regulatory mechanisms in aortic valve development and disease. Arteriosclerosis, thrombosis, and vascular biology, 34(4), pp.737–741

